# Molecular evolution, diversity and adaptation of H7N9 influenza A viruses in China

**DOI:** 10.1101/155218

**Authors:** Jing Lu, Jayna Raghwani, Jie Wu, Rhys Pryce, Thomas A. Bowden, Julien Thézé, Shanqian Huang, Yingchao Song, Lirong Zou, Lijun Liang, Ru Bai, Yi Jin, Pingping Zhou, Min Kang, Lina Yi, Oliver G. Pybus, Changwen Ke

## Abstract

A novel H7N9 avian influenza virus has caused five human epidemics in China since 2013. The substantial increase in prevalence and the emergence of antigenically divergent or highly pathogenic (HP) H7N9 strains during the current outbreak raises concerns about the epizootic-potential of these viruses. Here, we investigate the evolution and adaptation of H7N9 by combining publicly available data with newly generated virus sequences isolated in Guangdong between 2015-2017. Phylogenetic analyses show that currently-circulating H7N9 viruses belong to distinct lineages with differing spatial distributions. Using ancestral sequence reconstruction and structural modelling we have identified parallel amino-acid changes on multiple separate lineages. Furthermore, we infer mutations in HA primarily occur at sites involved in receptor-recognition and/or antigenicity. We also identify seven new HP strains, which likely emerged from viruses circulating in eastern Guangdong around March 2016 and is further associated with a high rate of adaptive molecular evolution.

## INTRODUCTION

Since its first detection in March 2013, avian influenza A virus (H7N9) has caused 1,452 human infections, resulting in 570 deaths (as of 9^th^ May 2017). Recurrent waves of human cases have been reported in 21 provinces of China, indicating sustained transmission of H7N9 viruses (Xiang et al., 2016). Moreover, since its emergence, H7N9 has reassorted with H9N2 avian influenza viruses that co-circulate in China, resulting in an increasingly diverse array of virus genomes (Cui et al., 2014, Lam et al., 2015, Wu et al., 2016). The current and fifth epidemic wave (2016/2017) has been marked by a notable increase in the number of human cases (677 between September 2016 and May 2017), making it the largest H7N9 virus outbreak since 2013. Moreover, the geographical distribution of human cases suggests that H7N9 is now more widespread, further raising public health concerns (Iuliano et al., 2017, Xiang et al., 2016).

Previous molecular surveillance studies have suggested that H7N9 epidemic lineages originate from two densely-populated areas, the Yangtze River Delta (YRD) region in eastern China, and the Pearl River Delta (PRD) region in central Guangdong (Wang et al., 2016). Preliminary epidemiological data suggests that the majority of human infections in the current, fifth, wave are caused by viruses belonging to the YRD lineage (Iuliano et al., 2017). Notably, in contrast to PRD viruses, viruses of the YRD lineage appear to exhibit reduced cross-reactivity with existing candidate vaccine virus (CVV) strains (Zhu et al., 2017). Furthermore, a subset of YRD lineage isolates have additionally acquired a highly pathogenic phenotype (Iuliano et al., 2017).

These observations suggest that the increased epidemic activity of H7N9 in China may be driven, at least in part, by ongoing viral evolution and adaptation. The decreased cross-reactivity and increased pathogenicity of some H7N9 viruses was discovered only very recently (Zhu et al., 2017), and the genetic diversity and evolution of the current fifth wave of H7N9 is not yet well understood. Key facts that urgently need to be established include (i) the geographic distribution of currently-circulating H7N9 lineages, (ii) the origin and genetic composition of newly-emerged highly-pathogenic H7N9 viruses and (iii) the evolutionary and structural characterisation of mutations associated with fifth-wave H7N9 viruses. In this study, we report 47 haemaglutinnin (HA) and 43 neuraminidase (NA) gene sequences of human- and poultry- derived H7N9 viruses that were isolated between 2015-2017 in Guangdong, China. By applying a structural and evolutionary analyses to these strains we characterise the evolution and emergence of currently-circulating H7N9 viruses in China.

## RESULTS

### Molecular epidemiology of H7N9 virus 2013-2017

Our analysis included 737 HA and 610 NA sequences from H7N9 viruses sampled between 2013 and 2017. As Figure 1a shows, the H7N9 epidemic lineage is geographically structured and classified into two major lineages, YRD and PRD. H7N9 has evolved in a ‘clock-like’ manner, i.e. there is a strong linear relationship between genetic divergence and sampling time (Figure 1b; correlation coefficient = 0.95). The estimated time of the most recent common ancestor (TMRCA) of the H7N9 HA sequences is November 2012 (95% credible region, CR = Oct 2012-Dec 2012). The corresponding molecular clock phylogeny for NA (Figure S1) also exhibits the YRD and PRD lineages, and has a similar estimated TMRCA, of September 2012 (95% CR = Jul 2012-Oct 2012). However, the topology of the NA phylogeny differs from that of HA, indicating reassortment between HA and NA during the emergence of the H7N9 epidemic lineage (Figures 1a and S1).

**Figure 1.**
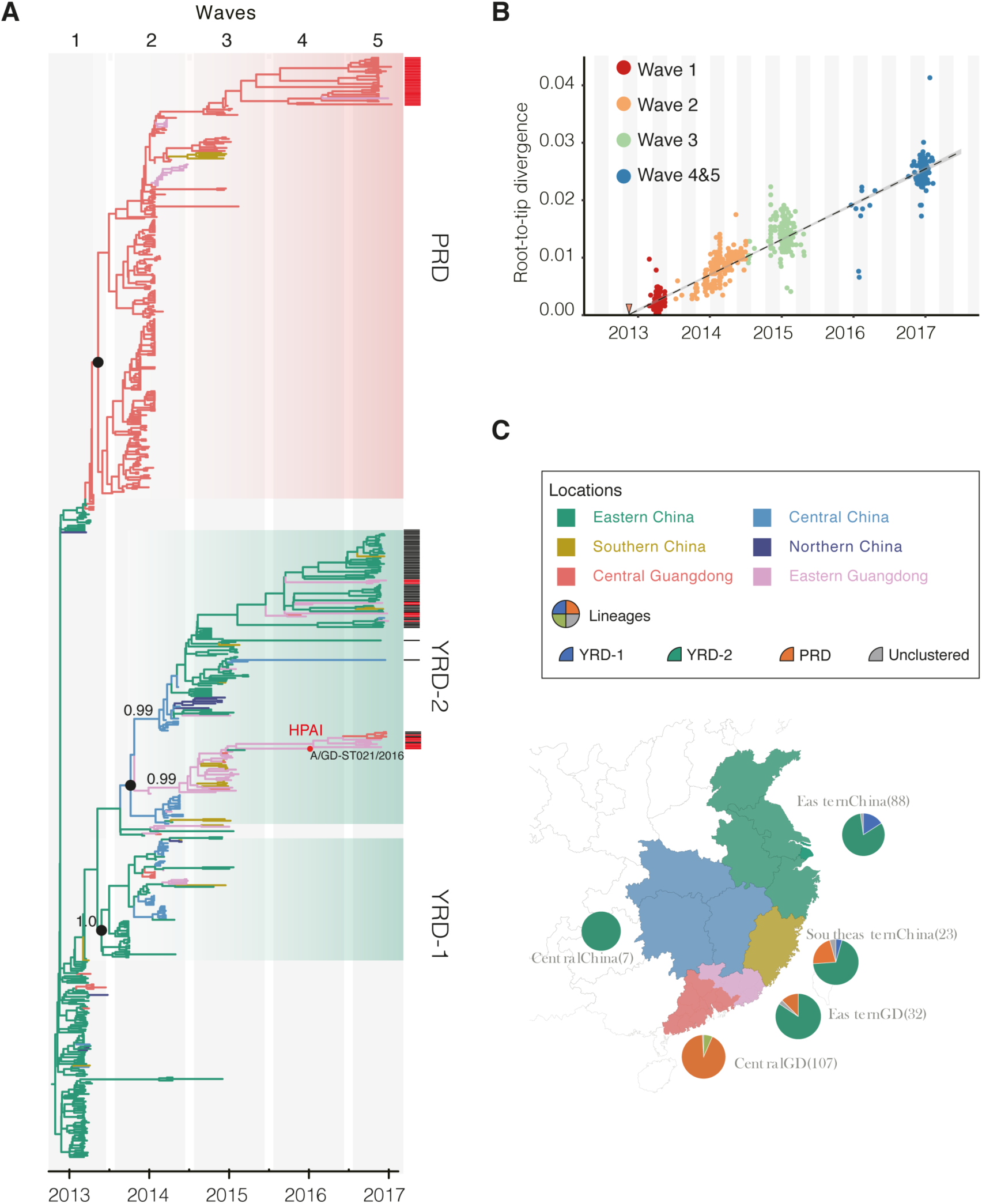
Genetic evolution and spatial spread of the H7N9 epidemic lineage in China 2013-2017. (a) Bayesian Maximum Clade Credibility (MCC) tree of HA gene from H7N9 viruses. Published sequences (from waves 4 and 5) from other studies are highlighted by black bars (Xiang et al., 2016, Iuliano et al., 2017) to the right of the tree; sequences reported in this study from Guangdong are highlighted by red bars. Branch colours represent the most probable ancestral locations of each branch. The theme colour is the same as shown in the right map. Two major lineages of H7N9 viruses are observed, denoted PRD and YRD. Black circles indicate posterior support >0.95. Location posterior support is shown for selected clades. A H7N9 strain closely related with HP H7N9 cluster is highlighted with red circle (b) A regression of root-tip divergence estimated from HA. The TMRCA of the H7N9 epidemic lineage is indicated by the red triangle (c) The geographical location and lineage classification of 374 H7N9 human viruses. The number of sequenced H7N9 viruses from each region are shown in braket. For visual clarity, sequences from Xinjiang Province in Northern China are not shown.

The YRD and PRD lineages are associated with different epidemiological patterns (Figures 1a and 1c). Specifically, the majority of PRD viruses (86%, 32/37) that were isolated during the fourth and fifth epidemic waves descended from viruses circulating in central Guangdong during earlier epidemic seasons (Figure 1a). In addition, PRD viruses isolated from the fourth and fifth waves are almost exclusively restricted to central (rather than eastern) Guangdong (Figures 1a and 1c). By contrast, YRD lineage viruses, which comprise two distinct sub-lineages (YRD-1 and YRD-2), have been exported to and become dominant in multiple regions as the epidemic has progressed (Lam et al., 2015), which indicates a comparatively broader geographic dissemination (Figure 1 and S1).

Our new isolates from eastern Guangdong, combined with isolates from eastern China (Xiang et al., 2016, Iuliano et al., 2017) suggest that recent YRD virus activity is primarily driven by the YRD-2 sublineage (Figure 1a). Whilst the YRD-1 sublineage has not been observed since the 2014-2015 epidemic season, the YRD-2 sublineage has increased in prevalence since its initial detection, in poultry samples from Jiangxi province in April 2014 (Figure 2A). The estimated TMRCA of the YRD-2 sublineage is December 2013 (95% HPD: Oct 2013 – Jan 2014). Importantly the frequency of YRD-2 virus detection among human cases has risen from 4% (4/96) in the second wave, to 70% (62/88) and 87% (71/82) in the third and fourth/fifth waves, respectively. In recent epidemic seasons, the YRD-2 sublineage has been associated with large outbreaks in eastern China and accounts for the majority of human cases outside central Guangdong (Iuliano et al., 2017).

**Figure 2.**
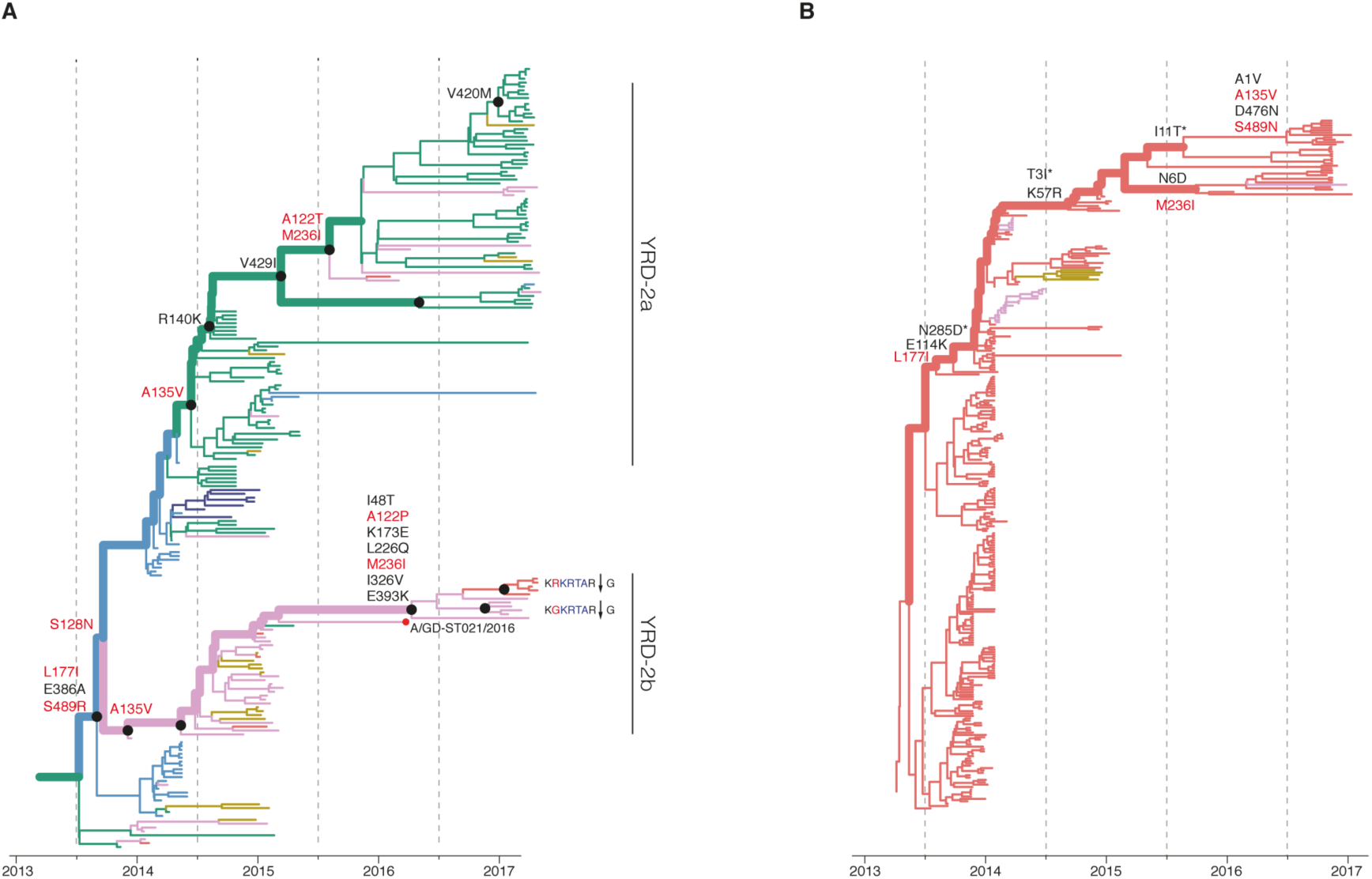
Reconstruction of amino acid changes along the trunk lineages of YRD-2 (A) and PRD (B) H7N9 viruses. The MCC tree of HA gene sequences from the YRD-2 lineage is shown. Branches are colored according to geographic locations, as in Figure 1. The thicker line indicates the trunk lineage leading up to the current fifth wave. Amino-acid changes along the trunk are shown. Sites undergoing parallel amino acid changes across multiple lineages are highlighted in red. Mutations correspond to H3 numbering scheme. Mutation sites which do not exist in H3 protein are marked by * are numbered according to H7 numbering.

Within YRD-2 we observed two clades (Figure 2a). The larger of these clades (YRD-2a) mainly circulates in central and eastern China, whilst the smaller (YRD-2b) is predominantly found in eastern Guangdong. YRD-2b also includes the recently identified highly pathogenic (HP) viruses (Figures 1 and 2a). To investigate these HP viruses, we undertook retrospective screening of poultry-related samples collected in Guangdong between January 2016 and February 2017, and identified seven HP influenza virus isolates that belong to the HP cluster within YRD-2b (Figure 1a). These HP viruses also form a distinct cluster within YRD-2 in the NA phylogeny (Figures S1). The most closely related isolate to the HP cluster is A/GD-ST021/2016, a strain isolated from a patient in eastern Guangdong (Figure 2a). Taken together, our analyses indicate that the HP clade likely emerged from YRD-2 viruses circulating in eastern Guangdong in 2016.

### Adaptive evolution in the YRD-2 virus lineage

Next we investigated whether the increasing prevalence of the YRD-2 sublineage might be associated with viral adaptation. We combined ancestral sequence reconstruction of YRD-2 and PRD HA gene sequences (Figure 2) by mapping residues that have undergone changes onto the crystal structure of the trimeric hemagglutinin. Our analysis reveals several notable amino-acid substitutions that occurred along the internal branches of the YRD-2 sublineage (Figure 2a).

Around the time of the second epidemic wave, ancestral YRD-2 viruses acquired several HA gene amino-acid changes, specifically L177I, G386A, S489R and S128N (based on H3 sequence numbering). Interestingly, three of these mutations (G386A, S489R, S128N) are located in solvent-accessible regions of HA (Figure 3; Table S2). Furthermore, we note that S128N occurs within the 130 loop and is proximal to the receptor surface (at approximately 20 Å distance); Figure 3).

**Figure 3.**
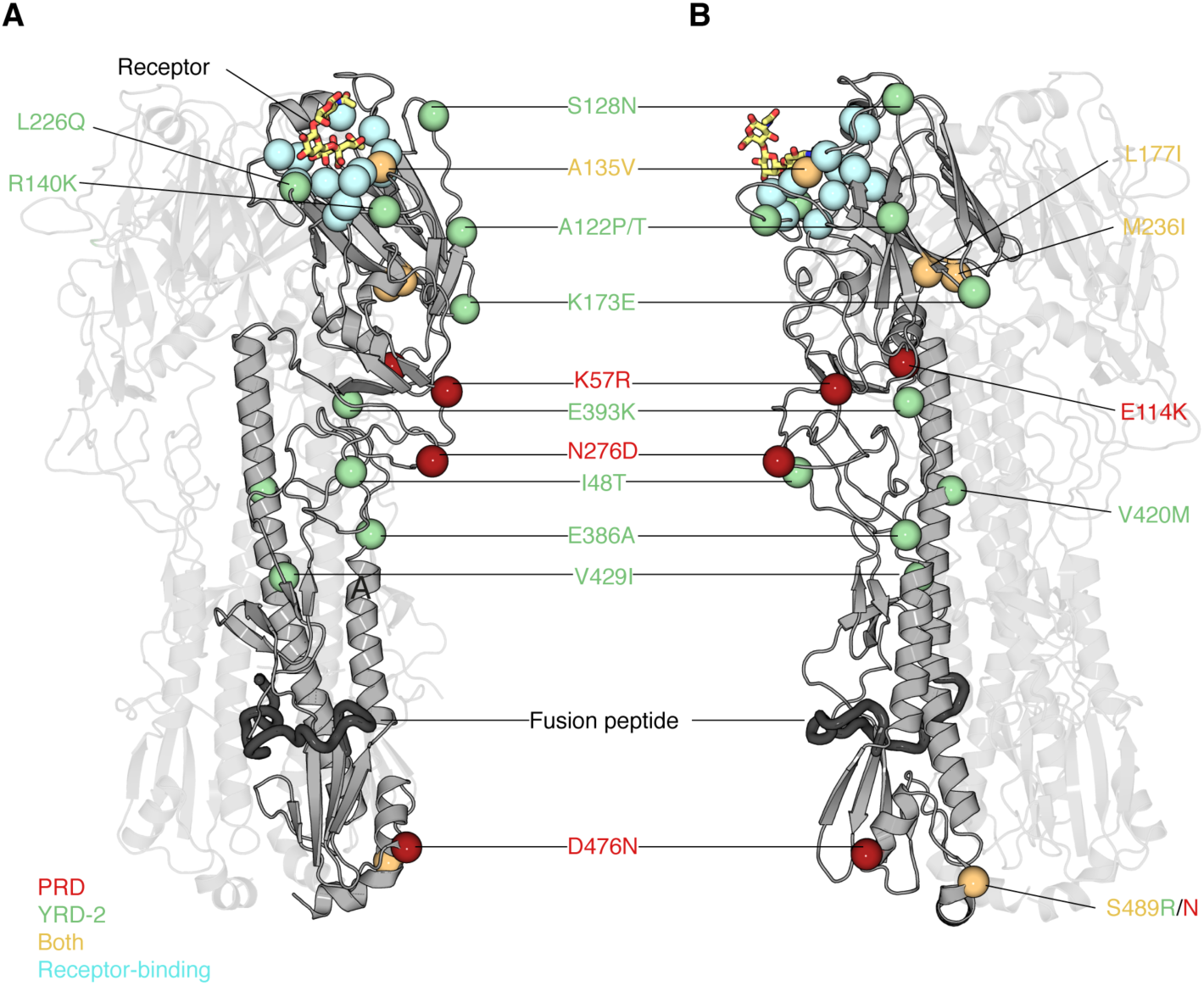
Structural analysis of amino acid changes within hemagglutinin in PRD and YRD-2 H7N9 viruses. (A) The crystal structure of the homotrimeric H7 hemagglutinin bound to a human receptor analogue (PDB ID, 4BSE (Xiong et al., 2013)) is shown in cartoon representation, and rotated 90° anticlockwise in panel (B). Two of the three protomers are displayed with high transparency to aid visualization. The carbon Ca positions of salient features are displayed as spheres: receptor binding residues are colored in cyan, mutations observed in the PRD and YRD-2 lineages are respectively represented by red and green residues. Residues that have experience mutations on both PRD and YRD-2 lineages are indicated in orange. The human receptor analogue a2,6-SLN is displayed as sticks colored according to constituent elements: carbon yellow, oxygen red, and nitrogen blue. The putative fusion peptide (Cross et al., 2009) is displayed as a thicker ‘cartoon’ and colored dark grey. Residues are numbered according to the H3 numbering system; further details about these mutations can be found in Table S2. A135 and L226 participate in receptor binding and thus are likely to modulate receptor specificity.

A highly notable result from our evolutionary analysis is that several HA sites acquired amino acid mutations *independently* in different phylogenetic clades. Firstly, four mutations (A135V, L177I, M236I and S489N) occurred independently along the trunk lineages that gave rise to current PRD and YRD-2 viruses (Figure 2b). Three of these (A135V, M236I and S489N) are observed only in the 4^th^ and 5th epidemic waves of the PRD lineage (Figure 2b). Secondly, comparison of the YRD-2a and YRD-2b clades also reveals parallel amino acid changes *within* the YRD lineage (A135V, M236I, and A122T/P; Figure 2a).

The observation of parallel amino-acid changes along those particular lineages that have persisted until the fifth epidemic wave (i.e. parallel changes between PRD and YRD-2 and between YRD-2a and YRD-2b) is suggestive of convergent, adaptive molecular evolution. Furthermore, there is some evidence that these changes have occurred in the same order, with mutation A135V preceding M236I, and A122T/P in both YRD-2a and YRD-2b (although not in PRD). All three parallel changes in YRD (A135V, M236I, A122T/P) are estimated to be fully or partially solvent accessible and the A135V mutation is located at the receptor-binding site (Figure 3). One subclade of the PRD lineage appears to have acquired the mutations A135V and S489N within the last 12 months (Figure 2b), and therefore we suggest this subclade should be closely monitored in the future.

In addition to the abovementioned parallel mutations, several amino acid changes are unique to each clade. Mutations that are specific to the trunk lineages within YRD-2a are R140K and V429I. While the solvent-inaccessibility and physicochemical similarity of the V429I mutation is unlikely to have a significant functional role, R140K occurs at a solvent accessible site known to be important for defining antigenicity and the substitution may alter the steric nature of the epitope (Wiley et al., 1981).

Interestingly, within the YRD-2b clade, HA acquired seven amino-acid changes (I48T, A122P, K173E, L226Q, M236I, I326V, and E393K) on the internal branch immediately ancestral to the HP cluster. This internal branch represents a period of approximately one year (see Figure 2a). While all of these changes appear in residues with partial or full solvent accessibility, mutations K173E, L226Q, and I326V are particularly noteworthy as they have arisen at or near known antigenic, receptor-binding, and proteolytic cleavage sites, respectively (Figure 3). Furthermore, these mutations in the HP cluster also coincide with the appearance of a four amino-acid insertion (KRTA) near the HA1-HA2 proteolytic cleavage site (Fig. 2a). Although we observed fewer amino acid changes in NA than in HA, we identified four amino-acid substitutions (A21T, V205I, A358D, and R430K; N2 numbering) along the branch that leads to the HP cluster (Figure S1; Table S3). Sites 358 and 430 are, fully and partially accessible, respectively. Site 21 was unobserved in the N9 crystal structure. Mutations at site 430 have been shown previously to contribute to antibody escape and to potentially affect NA structure (Lee and Air, 2002, Webster et al., 1987).

Lastly, we explored whether amino-acid changes in the HA gene during the H7N9 emergence were driven by positive selection. We find evidence for adaptive evolution in both the HA genes of PRD and YRD-1 lineages. We estimate the PRD lineage adapted at a rate of 0.80 (interquartile range, IQR = 0.21-0.95) adaptive substitutions per codon per year, and the YRD-1 lineage at a rate of 0.60 (IQR = 0.10-1.18). Within YRD-2, the estimated adaptation rate of the YRD-2a sublineage is ~1.84 (IQR: 1.09-2.14) adaptive substitutions per year, while that for the YRD-2b sublineage (which includes the HP cluster) is 3.12 (IQR: 2.45-3.79). These results indicate molecular adaptation across the whole H7N9 lineage and suggest that adaptation is faster in the two YRD-2 clades than in PRD and YRD-1.

## DISCUSSION

The results reported here show that H7N9 viruses of the YRD-2 lineage, which have been prevalent in the recent, fifth epidemic wave, comprise two geographically distinct clades (YRD2a and 2b) that have undergone substantial adaptive evolution. These clades are found primarily in eastern and central China and eastern Guangdong, respectively, and appear to have circulated in bird populations for about three years. Crucially, our ancestral state reconstruction analysis provides evidence that three successful lineages of H7N9 (PRD YRD-2a and 2b) have experienced multiple parallel amino acid changes (Fig. 2), suggesting the possible action of convergent molecular evolution. Lastly, we observe a higher rate viral adaptation in the eastern Guangdong YRD-2b clade compared to YRD-2a. This is concerning from a public health perspective, as HP H7N9 viruses have recently emerged within the YRD-2b clade. The introduction of HP avian influenza into domestic poultry can constitute a serious risk, as demonstrated by the emergence of goose-Guangdong lineage H5N1 HP viruses, which spilled back into wild birds and caused the longest global outbreak of HP avian influenza to date (Subbarao et al., 1998).

The parallel amino changes in clades YRD-2a and YRD-2b occur at three sites in HA (122, 135, and 236) that are located near or at known antigenic sites (Figure 3). The same mutations at the latter two sites are also observed in the PRD lineage (Figure 2). Taken together, these results suggesting adaptive convergent molecular evolution. Site 135 is located in the receptor-binding region and is in close proximity to antigenic site A, as defined by Wiley et al. (1981), so the observed A135V mutation in all three lineages may both modulate receptor affinity and contribute to immune escape (Figure 3). Interestingly, substitutions at site 135 have been suggested to play a role in viral antigenicity of HP viruses in the H7N1 and H7N7 outbreaks that occurred in Italy and the Netherlands, respectively (Monne et al., 2014, de Wit et al., 2010). Specifically, experimental studies indicate that the presence of threonine at position 135 in the HP H7N7 virus A/Netherlands/219/2003 confers broad-scale resistance to neutralizing monoclonal antibodies (mAbs) of the earliest strain of H7N9 virus (A/Shanghai/02/2013). Although the antigenic impact of the A135V mutation in contemporary (5^th^ wave) H7N9 viruses is currently unknown, its appearance on trunk branches in the YRD-2a, YRD-2b, and PRD lineages suggests that it is not directly linked with high pathogenicity (HP viruses are found only in clade YRD-2b). Nevertheless, amino acid variation at site 135 should be monitored in future H7N9 epidemic waves (Figure 2b). Amino acid changes at sites 122 and 236, which are located near known antigenic sites, appear to co-occur in both YRD-2 clades, and a parallel mutation at site 236 is also observed in the PRD lineage. While additional viral sequences will be necessary to resolve the relative timings of these mutations with certainty, especially in the YRD-2b clade, the possible functional dependence among changes these sites is intriguing, and they are worthwhile candidates for further experimental study. In particular, experiments to characterise the antigenicity of substitutions at 122, 135 and 236 sites are important for the selection of candidate vaccine virus strains.

We observed four other amino acid changes along the internal branches of the YRD-2 lineage. Two of these, S128N and L177I, are located near distinct antigenic sites that have been shown experimentally to reduce antibody responses when introduced together with A135T mutation in wild-type H7N9 viruses (Liu et al., 2016). Furthermore, site 128 in H5N1 viruses is known to play a role in viral attachment and mutation of this residue, due to its location in the 130-loop, has the potential to change virus preference for sialic-acid receptors with α2,3 or α2,6 linkage (Yamada et al., 2006). Although we do not assess the phenotypic effect of the S128N change in our work, it would be worthwhile investigating whether it has a role in effective mammalian transmission. While the other two substitutions occur in fully exposed residues in the HA2 domain (E386A and S489R), it is unclear whether these directly affect HA function. Since these sites are solvent accessible, they have the potential to be important for antigenicity.

Other amino acid changes in the YRD-2 lineage also occur in regions of HA that may affect viral antigenicity and/or host receptor binding. For example, the R140K change in the YRD-2a clade is located in antigenic site A, and has been observed in viruses isolated from ferrets experimentally infected with avian H7N9 viruses and has been linked to the antigenic drift of H5N1 viruses (Belser et al., 2013, Cattoli et al., 2011, Xu et al., 2014). A recent study has suggested that HP H7N9 viruses exhibit dual receptor binding properties, with an increased binding preference for avian receptor (possibly resulting from the reversion to glutamine at site 226) and a comparable binding preference for human receptor, when compared with Anhui/1/2013 (Zhu et al., 2017).

In the recent epidemic waves, YRD-2 H7N9 viruses were dominant in four of the five regions that were most affected by H7N9 (Figure 1). In this study we identified several mutations along the trunk of YRD-2 lineage that have greatly increased in frequency in human viruses, or become fixed in the virus population (Figure S2). Although PRD viruses are mainly observed in central Guangdong and have lower adaptation rate than YRD-2 viruses, they too appear to have fixed a number of mutations between the 3^rd^ and 5^th^ epidemic waves.

H7N9 virus epidemics are currently restricted to China, except for a few sporadic exported human cases (Xiang et al., 2016), and H7N9 viruses do not appear to be able transmit directly among humans. However, recent work suggests that the newly-emerged HP H7N9 viruses have higher infectiousness in poultry than related low pathogenicity strains (Zhu et al., 2017) and these HP viruses are associated with a high inferred rate of molecular adaptation. This raises the possibility of global dissemination of H7N9 through migration of wild birds, in a manner similar to that observed for HP H5N1 viruses first identified in Guangdong (Subbarao et al., 1998). Recent surveillance indicates that HP viruses have already spread to several provinces and are responsible for large outbreaks in central and northern China with high poultry mortality (<u>www.moa.gov.cn</u>). Even though HP H7N9 infection in wild birds remains largely unknown, close monitoring of H7N9 viruses in neighboring countries, including Japan, Vietnam, Thailand and Indonesia, is warranted.

## Author Contributions

C.K., J.W., J. L designed the study. J.L., Y.S., L.Z., L.L., R.B., Y.J., P.Z., M. K. and L.Y. prepared sample collection and genome sequencing. J. L., J.R., R.P., T.B., J.T., S.H. and O.P. analyzed the data. J. L., J.R., R.P., T.B., and O.P. interpreted the data. J. L., J.R., R.P., T.B., and O.P. prepared the figures. J. L., J.R., R.P., T.B., and O.P. wrote the article.

## Acknowledgements

This work was supported by grants from the National Natural Science Foundation of China (81501754) and the National Key Research and Development Program of China (2016YFC1200201). We thank the MRC (MR/L009528/1 to T.A.B.). The Wellcome Trust Centre for Human Genetics is supported by Wellcome Trust Centre grant 203141/Z/16/Z.

## STAR METHODS

### Sample Collection

Samples from suspected human cases were initially tested for avian influenza A virus in provincial clinics in Guangdong, China. Specimens with positive results were subsequently tested in prefecture-level CDC laboratories for subtypes H5, H7, and H9, as previously described (Ke et al., 2014, Lu et al., 2014). Positive specimens were sent to Guangdong Provincial CDC and confirmed by H7N9-NA RT-PCR. For poultry-related samples, we obtained samples from locations where poultry were housed and processed (e.g. cages, feeding-troughs, de-feathering machines). For each of the 21 prefectural-level divisions of Guangdong province, Guangdong CDC tested 10 wet-swab poultry specimens per month, as previously described (Ke et al., 2014). When a human H7N9 case was confirmed at a given location, and the patient had exposure to a specific live poultry market (LPM), at least 20 environmental samples were collected from that market. Respiratory specimens were collected from suspected H7N9 cases by the Ministry of Health, P. R. of China. Written consent was obtained from each patient or their guardian(s) when samples were collected.

Following RT-PCR testing, H7N9 positive swabs and sputum samples were blindly passaged for 2 to 3 generations in 9-10 day old embryonated chicken eggs. Hemagglutination-positive allantoic fluids were collected, and viruses were subtyped by hemagglutination and neuraminidase inhibition assays using a panel of reference antiserum, as described in (Huang et al., 2012). H7 and N9 genes segments of the selected isolates were amplified as previously described (Wu et al., 2016).

### Sequence alignment

In this study, we sequenced 47 HA and 41 NA sequences from 20 human cases and 28 poultry-related samples, all belonging to the fourth and fifth epidemic waves of H7N9 (GISAID accession numbers: EPI866538-866577, 972231-972236, 972238-972303, 974029, 974523, 974539-974542, 997159-997160). These new H7N9 sequences were combined with all publicly available H7N9 gene sequences whose sampling dates and locations were known. Only one representative was retained when very similar sequences (nucleotide identity >99%) were sampled from the same location and time. Two gene sequence datasets were generated: H7 HA (*n*=737) and N9 NA (*n*=610). Multiple sequence alignments were constructed using ClustalW (Larkin et al., 2007) and then edited by hand using AliView (Larsson, 2014).

### Molecular clock phylogenetic analysis

Maximum-likelihood (ML) phylogenies were estimated from the H7 and N9 alignments using RAxML (Stamatakis, 2006), under the GTR+Γ nucleotide substitution model (Yang, 1994). For each alignment, accumulation of genetic divergence through time was assessed from mid-point rooted ML phylogenies using the regression approach implemented in TempEst (Rambaut et al., 2016).

Molecular clock phylogenies were estimated using the Bayesian MCMC approach implemented in BEAST v 1.8 (Drummond et al., 2012). Specifically, we employed a SRD06 nucleotide substitution model (Shapiro et al., 2006), a uncorrelated log-normal relaxed molecular clock model (Drummond et al., 2006), and a Bayesian Skygrid coalescent model (Gill et al., 2013). Four independent MCMC runs of 1.5×10^8^ steps were computed for each alignment. A subset of 2000 phylogenies was extracted from the posterior tree distribution and subsequently used as an empirical tree distribution for phylogeographic analyses (Lemey et al., 2014). Maximum clade credibility (MCC) trees for each alignment were computed using TreeAnnotator (Drummond et al., 2012).

### Phylogeographic analysis of the H7N9 viral epidemic

We used the discrete phylogeographic method (Lemey et al., 2009), implemented in BEAST, to investigate the spatial dynamics of H7N9 lineages among six regions in China: (i) Eastern China (Anhui, Shanghai, Zhejiang, Jiangsu and Shandong), (ii) Central China (Jiangxi and Hunan), (iii) Northern China (Beijing, Henan, Hebei, Xinjiang), (iv) Southeastern China (Fujian), (v) Central Guangdong (Guangzhou, Huizhou, Foshan, Dongguan, Zhongshan, Shenzhen, Jiangmen, Zhaoqing Yangjiang, Maoming and Yunfu), and (vi) Eastern Guangdong (Meizhou, Heyuan, Chaozhou, Jieyang, Shantou, Shanwei and Shaoguan). Sporadic human cases detected in Malaysia and Taiwan are thought to be exports from China, hence available epidemiological information was used to assigned their location to the most likely source location in China. Hong Kong and Central Guangdong were treated as a single location due to their proximity. The number of reported H7N9 infections and viral sequences used are provided in Table S1. To estimate the directionality of virus lineage movement, we employed an asymmetric continuous-time Markov chain (CTMC) phylogeographic model (Edwards et al., 2011) together with a Bayesian stochastic search variable selection (BSSVS) procedure (Lemey et al., 2009).

### Inferring the phylogenetic distribution of amino acid changes

The phylogenetic positions of amino-acid changes among H7N9 isolates were investigated using the HA and NA MCC trees. The maximum posterior probability amino acid sequences for each internal node were estimated using BEAST, under a JTT amino-acid substitution model (Jones et al., 1992), gamma-distributed among-site rate heterogeneity (Yang, 1996) and a strict molecular clock model. To infer amino acid substitutions along the ‘trunk branches of the H7N9 phylogeny, amino-acid changes were mapped onto internal branches using a Java script (available on requested). ‘Trunk branches’ were those that subtended more than 5 terminal nodes in the fifth epidemic wave.

### Structure-based mapping analysis

We used the crystal structure of the HA (Protein Data Bank ID 4BSE: (Xiong et al., 2013)) and NA (Protein Data Bank ID 2C4L) glycoproteins from a H7N9 influenza viruses to map the amino acid changes identified by the abovementioned evolutionary analysis. Residue mapping onto the H7 and N9 structures was performed using PyMol (Schrödinger, 2015). Solvent accessibility was calculated for trimeric hemagglutinin, with the ligands removed, using ESPript (Gouet et al., 1999), and receptor binding residues were determined using ‘CONTACT’ in CCP4 (Winn et al., 2011).

### Estimating rates of viral molecular adaptation

Rates of adaptive substitution in H7N9 HA and NA genes were estimated using an established population genetic method related to the McDonald-Kreitman test (Bhatt et al., 2011, Raghwani et al., 2016). A consensus of H7N9 first wave sequences was used as an ‘outgroup’ to compute the derived and ancestral mutational site frequencies in each subsequent wave. Specifically, polymorphisms were classified into three categories (low, intermediate, and high) according to their frequency in the population (0-15%, 15%-75%, and 75%-100%, respectively). The number of adaptive substitutions was calculated from the number of synonymous and non-synonymous sites in each category and statistical uncertainty was assessed using a bootstrap approach (1000 replicates). See Bhatt et al. (2011), Raghwani et al. (2016) for a detailed description.

